# The actin depolymerizing factor StADF2 alters StREM1.3 plasma membrane nanodomains to inhibit the *Potato Virus X*

**DOI:** 10.1101/2023.01.25.525625

**Authors:** Marie-Dominique Jolivet, Paul Gouguet, Anthony Legrand, Kaltra Xhelilaj, Natalie Faiss, Aurélie Massoni-Laporte, Terezinha Robbe, Isabelle Sagot, Marie Boudsocq, Sylvie German-Retana, Suayib Üstün, Antoine Loquet, Birgit Habenstein, Véronique Germain, Sébastien Mongrand, Julien Gronnier

**Author notes:** These authors (M-D. J, P.G.) contributed equally to the work and should be considered as co-first authors. These authors (V.G, S.M., J.G.) contributed equally to the work and should be considered as co-last authors. Corresponding author: Julien Gronnier < > Zentrum für Molekularbiologie der Pflanzen (ZMBP), Eberhard Karls Universität Tübingen, Auf der Morgenstelle 72076, Tübingen, Germany.

## Abstract

The dynamic regulation of the plasma membrane (PM) organization at the nanoscale emerged as a key element shaping the outcome of host-microbe interactions. Protein organization into nanodomains (ND) is often assumed to be linked to the activation of cellular processes. In contrast, we have previously shown that the phosphorylation of the *Solanum tuberosum* REM1.3 (StREM1.3) N-terminal domain disperses its native ND organization and promotes its inhibitory effect on *Potato Virus X* (PVX) cell-to-cell movement. Here, we show that the phosphorylation of StREM1.3 modify the chemical environment of numerous residues in its intrinsically-disordered N-terminal domain. We leveraged exploratory screens to identify potential phosphorylation-dependent interactors of StREM1.3. Herewith, we uncovered uncharacterized regulators of PVX cell-to-cell movement, linking StREM1.3 to autophagy, water channels and the actin cytoskeleton. We show that the *Solanum tuberosum* actin depolymerizing factors 2 (StADF2) alters StREM1.3 NDs and limits PVX cell-to-cell movement in a REMORIN-dependent manner. Mutating a conserved single residue reported to affect ADFs affinity to actin inhibits StADF2 effect on StREM1.3 ND organization and PVX cell-to-cell movement. These observations provide functional links between the organization of plant PM and the actin cytoskeleton and suggests that the alteration of StREM1.3 ND organization promotes plant anti-viral responses. We envision that analogous PM re-organization applies for additional signaling pathways in plants and in other organisms.

## INTRODUCTION

The plasma membrane (PM) actively hosts, modulates and coordinates a myriad of signaling events essential to the development and survival of all living organisms. Across the tree of life, PM lipids and proteins have been found to be dynamically organized into diverse nanoscopic environments termed nanodomains (NDs) (Gronnier et al., 2018; Jacobson et al., 2019; Lopez & Koch, 2017; Malinsky et al., 2013). NDs have been proposed to provide dedicated biochemical and biophysical environments to ensure acute, specific and robust signaling events (Kusumi et al., 2012; Sezgin et al., 2017). In plants, emerging evidence suggest that ND formation and integrity rely on the intimate interplay between lipids, proteins, the cell wall and the cytoskeleton (Galindo-Trigo et al., 2020; Gronnier et al., 2018, 2019; Jaillais & Ott, 2020). The molecular events underlying context-dependent PM re-organization and their functional consequences remain largely unknown.

REMORINs are structural components of the PM playing regulatory functions in plant-microbe interactions and hormone signaling (reviewed in Gouguet et al., 2021). REMORINs of different groups (Raffaele et al., 2007) tend to form distinct and coexisting NDs proposed to regulate specific signaling pathways (Bücherl et al., 2017; Jarsch et al., 2014). The molecular bases for REMORINs ND organization are best characterized for group 1 REMORINs (Gronnier et al., 2018; Jaillais & Ott, 2020). Group 1 REMORINs NDs are proposed to be liquid-ordered and enriched in sterols and saturated lipids (Demir et al., 2013; Gronnier et al., 2017; Mamode Cassim et al., 2019; Raffaele et al., 2009). Their native organization is dictated by direct interactions with anionic phospholipids (Gronnier et al., 2017; Legrand et al., 2019; Legrand et al., 2022; Perraki et al., 2012), and can be modulated by post-translational modifications such as S-acylation (Fu et al., 2018; Konrad et fal., 2014) and phosphorylation (Perraki et al., 2018). Group 1 REMORINs regulate cell-to-cell movement of viruses belonging to diverse genus (Cheng et al., 2020; Fu et al., 2018; Huang et al., 2019; Raffaele et al., 2009; Rocher et al., 2022). Notably, their function in hindering *Potexviruses*, to which the *Potato Virus X* (PVX) belongs, is conserved in *Nicotiana benthamiana, Solanum lycopersicum* and *Arabidopsis thaliana* (Abel et al., 2021; Perraki et al., 2018; Perraki et al., 2014; Raffaele et al., 2009), suggesting an ancestral function conserved across at least 100 million years of evolution. Most REMORINs bear a predicted intrinsically-disordered N-terminal domain found to be phosphorylated in multiple plant species and across a wide range of physiological conditions (Gouguet et al., 2021). Phosphorylation affects the chemistry, structure, and conformational ensemble of intrinsically-disordered domains (IDD), and consequently their association with protein partners (Mukhopadhyay et al., 2022; Wright & Dyson, 2014). In view of their PM ND organization, their ability to homo-oligomerize, and their putative IDD, REMORINs have been proposed to act as scaffolds. Here, we show that the N-terminal domain of StREM1.3 is intrinsically disordered and that its phosphorylation modifies the chemical environment of numerous residues. Leveraging yeast two-hybrid screens, we identified interactors of a phosphomimic variant of StREM1.3 and uncovered uncharacterized regulators of PVX infection. We show that the *Solanum tuberosum* actin depolymerizing factors 2 (StADF2) alters StREM1.3 NDs and limits PVX propagation in a REMORIN-dependent manner. We observed that StADF2 activity is affected by mutating a conserved residue regulating ADFs affinity to actin. Altogether, these observations suggest that active modulation of REMORIN ND organization promotes plant anti-viral responses.

## RESULTS

### StREM1.3 N-terminal domain is intrinsically disordered and its phosphorylation leads to changes in the chemical environment of numerous residues

We previously showed that phosphorylation of the *Solanum tuberosum* REMORIN1.3 (StREM1.3) N-terminal domain regulates its ND organization and function (Perraki et al., 2018). Notably, using photoactivated localization microscopy, we observed that a phosphomimic form of StREM1.3, StREM1.3^S74D T86D S91D^ (further referred to as StREM1.3^DDD^) which restricts PVX cell-to-cell movement, presented a dispersed organization (Perraki et al., 2018). We confirmed this observation using laser scanning confocal microscopy (Figure 1A). However, the molecular bases underlying StREM1.3 ND dispersion remain unknown. StREM1.3 N-terminal domain is predicted to be intrinsically disordered (Marín et al., 2012; Perraki et al., 2018). We used liquid-state NMR spectroscopy to characterize the properties of StREM1.3 N-terminal domain. ^1^H-^15^N HMQC spectra of untagged StREM1.3^1-116^ purified in native and denaturing conditions (Figure S1A) both displayed a narrow δ(1H) distribution (between ~ 7.5-8.5 ppm, Figure S1B), demonstrating that StREM1.3 N-terminal domain is intrinsically disordered.

**Figure 1.**
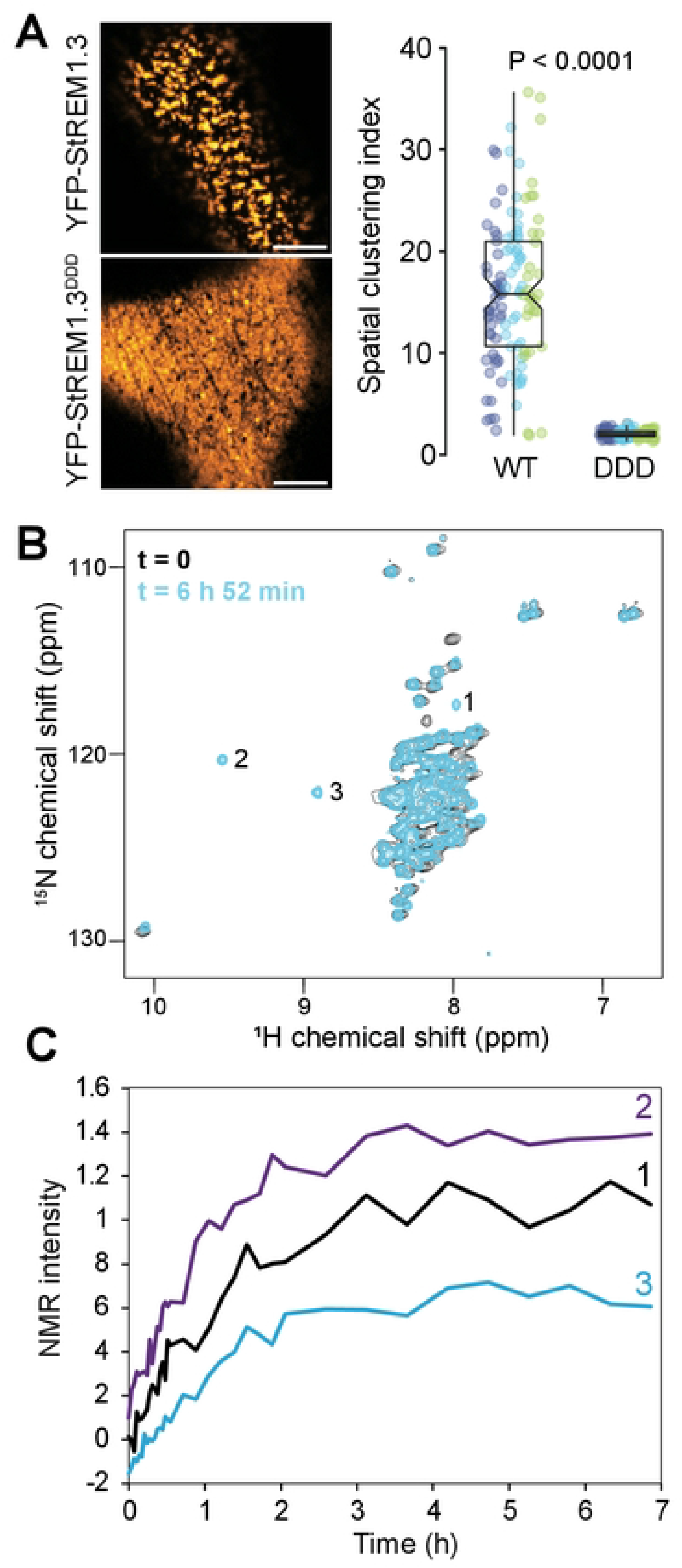
Phosphorylation modulates StREM1.3 intrinsically disordered domain. **A**. Representative confocal micrograph showing the surface of *N. benthamiana* leaf epidermal cells expressing YFP-StREM1.3^WT^ and YFP-StREM1.3^DDD^ (left) and corresponding quantification of YFP-StREM1.3^WT^ and YFP-StREM1.3^DDD^ spatial clustering index (right). Graphs are notched box plots, scattered data points show measurements, colors indicate independent experiments, n=38 cells for YFP-StREM1.3 and n=39 cells for YFP-StREM1.3^DDD^. P values report two-tailed non-parametric Mann-Whitney test. Scale bar indicates 5 μm. **B**. ^1^H-^15^N HMQC spectra of StREM^1-116^ before (black) and after (blue) addition of AtCPK3. Peaks labelled 1,2 and 3 appeared over time. Each spectrum was acquired using 2 scans for an experimental time of 1 min 42 s each. Temperature: 20°C. **C**. NMR intensities of peaks 1, 2 and 3 (Figure 1B) over time after the addition of AtCPK3.

We then asked how phosphorylation modulates StREM1.3 N-terminal domain. The *Arabidopsis thaliana* calcium-dependent protein kinase 3 (AtCPK3) was previously shown to phosphorylate group 1 REMORINs (Mehlmer et al., 2010; Perraki et al., 2018) and to restrict PVX cell-to-cell movement in a REMORIN-dependent manner (Perraki et al., 2018). Liquid-state NMR spectroscopy showed that the addition of AtCPK3, which phosphorylates StREM1.3 N-terminal domain *in vitro* (Legrand et al., 2022; Perraki et al., 2018) (Figure S2), induced chemical shift perturbations on several residues without altering the intrinsically disordered nature of StREM1.3^1-116^ (Figures 1B and 1C). The chemical shifts of the amide groups are very sensitive to the local chemical environment of the concerned residues. These observations indicate the modification of the chemical environment for multiple residues induced by phosphorylation which may modulate StREM1.3 intra- and intermolecular interactions. We therefore hypothesized that the phosphorylation of StREM1.3 intrinsically disordered N-terminal domain may modulate its association with specific protein partners to inhibit PVX cell-to-cell movement.

### A StREM1.3 phosphomimic-based yeast two hybrid exploratory screen identifies uncharacterized regulators of PVX cell-to-cell movement

To identify proteins which may associate with StREM1.3 in a phosphorylation-dependent manner, we used exploratory split-ubiquitin yeast two-hybrid (SUY2H) screens comparing WT and phosphomimic variant of StREM1.3, StREM1.3^DDD^. First, we tested whether StREM1.3 was a functional bait in SUY2H. We observed that expression of Cub-StREM1.3^WT^ or Cub-StREM1.3^DDD^ did not lead to autoactivation and that StREM1.3 self-association (Bariola et al., 2004.; Martinez et al., 2019; Perraki et al., 2014) could be resolved in this system, indicating that both Cub-StREM1.3 or Cub-StREM1.3^DDD^ were functional in SUY2H assays (Figure S3A and S3B). Further, we observed that GFP-tagged StREM1.3 and StREM1.3^DDD^ localized to the PM when expressed in yeast (Figure S3C). Interestingly, both proteins formed PM domains in yeast (Figure S3C), suggesting that StREM1.3^DDD^ dispersed organization as observed in *N. benthamiana* (Figure 1A) is conditioned by plant factors.

To enrich for relevant factors during viral infection, we generated a cDNA library using mRNA extracted from both PVX-infected and healthy *N. benthamiana* leaf epidermis (Figure S4A and S4B). We used peeled epidermis to limit the occurrence of chloroplastic proteins in our screen as previously reported (Bernard et al., 2012). We confirmed the presence of PVX:GFP in the infected epidermis by confocal microscopy (Figure S4C). Side-by-side screens of this library by SUY2H revealed an approximate 6-fold increase in the number of positive clones obtained with StREM1.3^DDD^ compared to StREM1.3 (Figures 2A, 2B and 2C), suggesting that the phosphomimic form of StREM1.3 has an increased ability to generate protein-protein interactions. Sequencing of the clones presenting reliable interaction identified 37 and 140 distinct proteins for StREM1.3 and StREM1.3^DDD^ respectively (Figure 2D and supplemental table 1). Among them, only 6 proteins were identified in both screens (Figure 2D) suggesting that StREM1.3 and StREM1.3^DDD^ have largely distinct interactomes. Among identified proteins are orthologs of Arabidopsis 14-3-3 and HYPERSENSITIVE INDUCED REACTION proteins which have been previously shown to associate with group 1 REMORINs (Huang et al., 2019; Lv et al., 2017), pointing toward the functional relevance of the identified proteins. To further explore the role of REM interactors in PVX infection, we prioritized candidates based on the number of clones identified in SUY2H screen, their co-purification and/or co-expression with Arabidopsis REMs in omics studies and cloned them as translational fusions with the HA epitope-tag for functional studies. Among the 10 candidates tested in PVX:GFP cell-to-cell propagation assays (Figures 3A and 3B), the expression of 7 candidates impaired PVX propagation: the Plasma membrane intrinsic protein 1C (NbPIP1C), Delta tonoplast integral protein (NbTIP2;1), Thioredoxin superfamily protein 4 (NbTRX4), Metallothionein 2B (NbMT2B), Calreticulin 3 (NbCRT3), Autophagy 8i (NbATG8i) and Actin Depolymerizing Factor 3 (NbADF3) (Figure 3B, Figure S5). Altogether, these observations both suggest that the phosphorylation of StREM1.3 promotes its association with specific proteins to limit PVX cell-to-cell movement and link plant response to PVX with the autophagy pathway and water channels notably.

**Figure 2.**
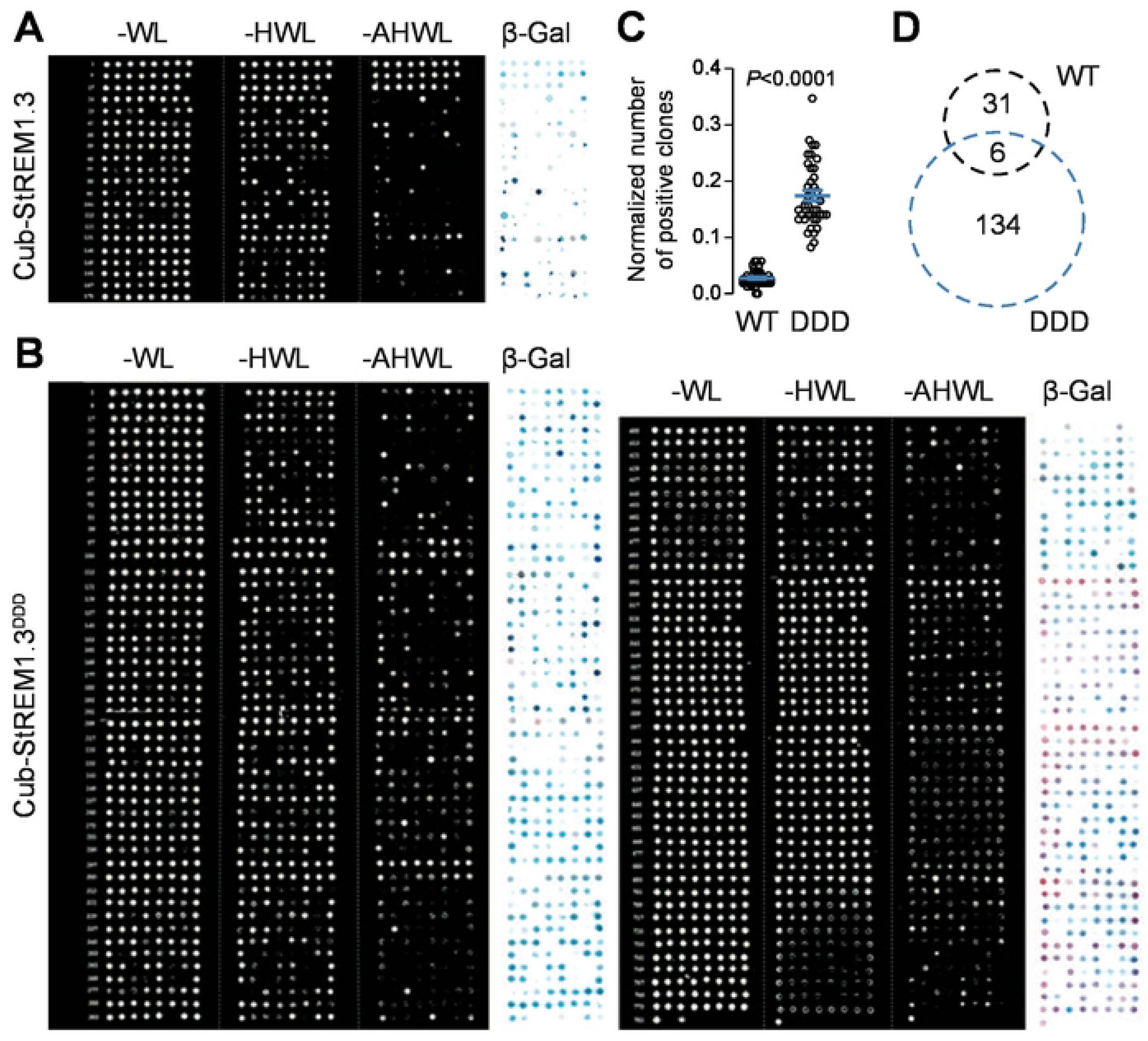
Comparative analysis of StREM1.3 and StREM1.3^DDD^ interactomes in SUY2H. **A-B**. Drop-tests of SUY2H assays performed with yeast clones retrieved from two independent side-by-side screens performed using Cub-StREM1.3 (**A**) or Cub-StREM1.3^DDD^ (**B**) as bait and constructs from the *N. benthamiana* leaf epidermis cDNA library. **C**. Quantification of the number of positive clones retrieved from Cub-StREM1.3 and Cub-StREM1.3^DDD^ screens, observed on -AHWL plates 4 days after transformations normalized by transformation efficiency for each screen (expressed in ‰ of theoretically tested clones). Data points show values obtained for individual -AHWL plates. P values report two-tailed non-parametric Mann-Whitney test. **D**. Venn diagram depicting the number of proteins identified with Cub-StREM1.3 and Cub-StREM1.3^DDD^.

**Figure 3.**
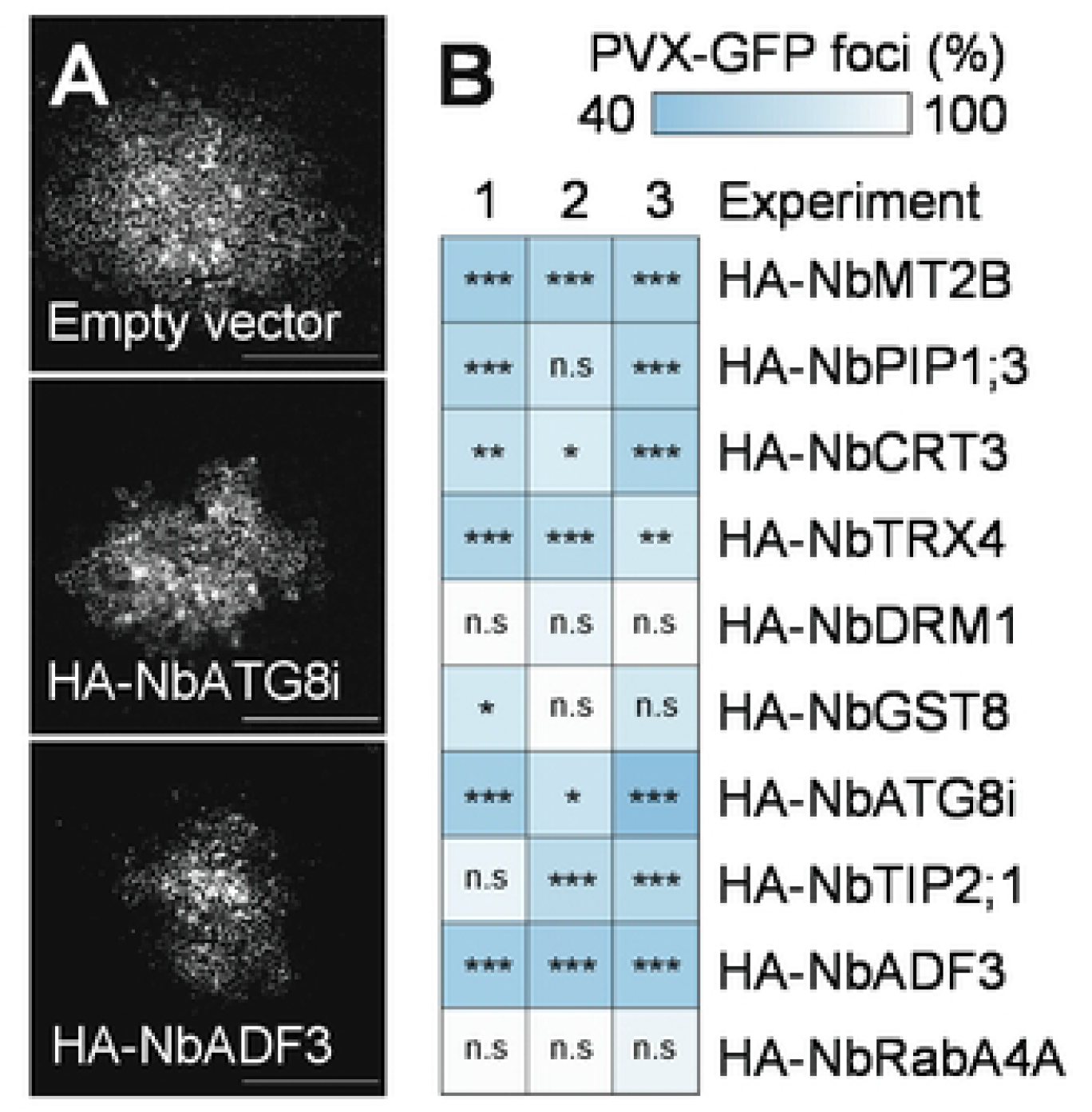
Functional analysis of StREM1.3’s putative interactors upon PVX infection. **A**. Representative images of PVX:GFP infection foci of *N. benthamiana* leaves expressing empty vector (control condition) and 2 selected HA-tagged candidates. Scale bar = 500 μm. **B**. Quantification of PVX infection assays. The table is color coded based on the mean PVX:GFP infection foci area observed upon the overexpression of individual candidates normalized to the mean PVX:GFP infection foci area of the corresponding empty vector condition. Each candidate was N-terminally-fused to an HA-tag and expressed under the control of a 35S promoter. Number of infection foci n=182 for HA-NbMT2B, n=219 for HA-NbPIP1;3, n=177 for HA-NbCRT3, n=249 for HA-NbTRX4, n=208 for HA-NbDRM1, n=223 for HA-NbGST8, n=114 for HA-ATG8i, n=228 for HA-TIP2;1, n=209 for HA-NbADF3, n=175 for HA-RabA4A. Stars report by Dunn’s comparisons test, p<0.05: * p<0.01: **, p<0.001: ***, non-significant (n.s): p>0.05.

### StADF2 affects StREM1.3 nanodomains and inhibits PVX cell-to-cell movement in a REMORIN-dependent manner

Among the genes tested, the *N. benthamiana* actin depolymerizing factor (ADF) 3 (NbADF3, NbS00025994g0001.1) was one of the most potent in limiting PVX cell-to-cell movement (Figure 3B). To corroborate the potential implication of an ADF-REMORIN module involved in response to PVX, we first tested the ability of an ortholog of NbADF3 from *Solanum tuberosum* (StADF2, Soltu.DM.04G007350) to associate with StREM1.3. Co-immunoprecipitation experiments, using transient expression in *N. benthamiana* leaves, showed that StADF2 and StREM1.3 associate *in planta* (Figure 4A). We next tested whether StADF2 limits PVX infection. Using group 1 *REMORINs* knock-down stable transgenic *N. benthamiana* lines (hpREM #1.4 and #10.2; Perraki et al., 2018), we observed that the transient overexpression of StADF2 limited PVX cell-to-cell movement in a REMORIN-dependent manner (Figure 4B, 4C). These results indicate that ADFs and REMORINs cooperate to limit PVX infection. Because the ND organization of the *Arabidopsis thaliana* REM1.2 and of the *Medicago truncatula* SYMBIOTIC REMORIN (SYMREM; MtREM2.2) rely on the integrity of the actin cytoskeleton (Liang et al., 2018; Szymanski et al., 2015) and that the active form of StREM1.3 correlates with a disperse PM organization (Figure 1A; Perraki et al., 2018), we wondered whether StADF2 could regulate StREM1.3 ND organization to inhibit PVX. We observed that overexpression of StADF2 affected YFP-StREM1.3 ND organization (Figures S6A and S6B), suggesting that StREM1.3 ND organization relies on actin cytoskeleton integrity. In good agreement, cytochalasin D treatment, which inhibits actin polymerization, was sufficient to affect YFP-StREM1.3 ND organization (Figures S6C and S6D). The affinity of ADFs for actin is decreased by their phosphorylation on a conserved N-terminally located Serine (Figure S7; Augustine et al., 2008; Dong & Hong, 2013; Y. J. Lu et al., 2020; Porter et al., 2012; Ressad et al., 1998). Interestingly, we observed that replacing the corresponding Serine residue by an Aspartic acid in StADF2 (StADF^S6D^) altered its ability to modify StREM1.3 ND organization (Figures 5A and 5B) and to restrict PVX cell-to-cell movement (Figure 5C and 5D). Altogether, these data indicate that StADF2 restricts PVX cell-to-cell movement by actively modulating actin cytoskeleton and REMORINs ND organization.

**Figure 4.**
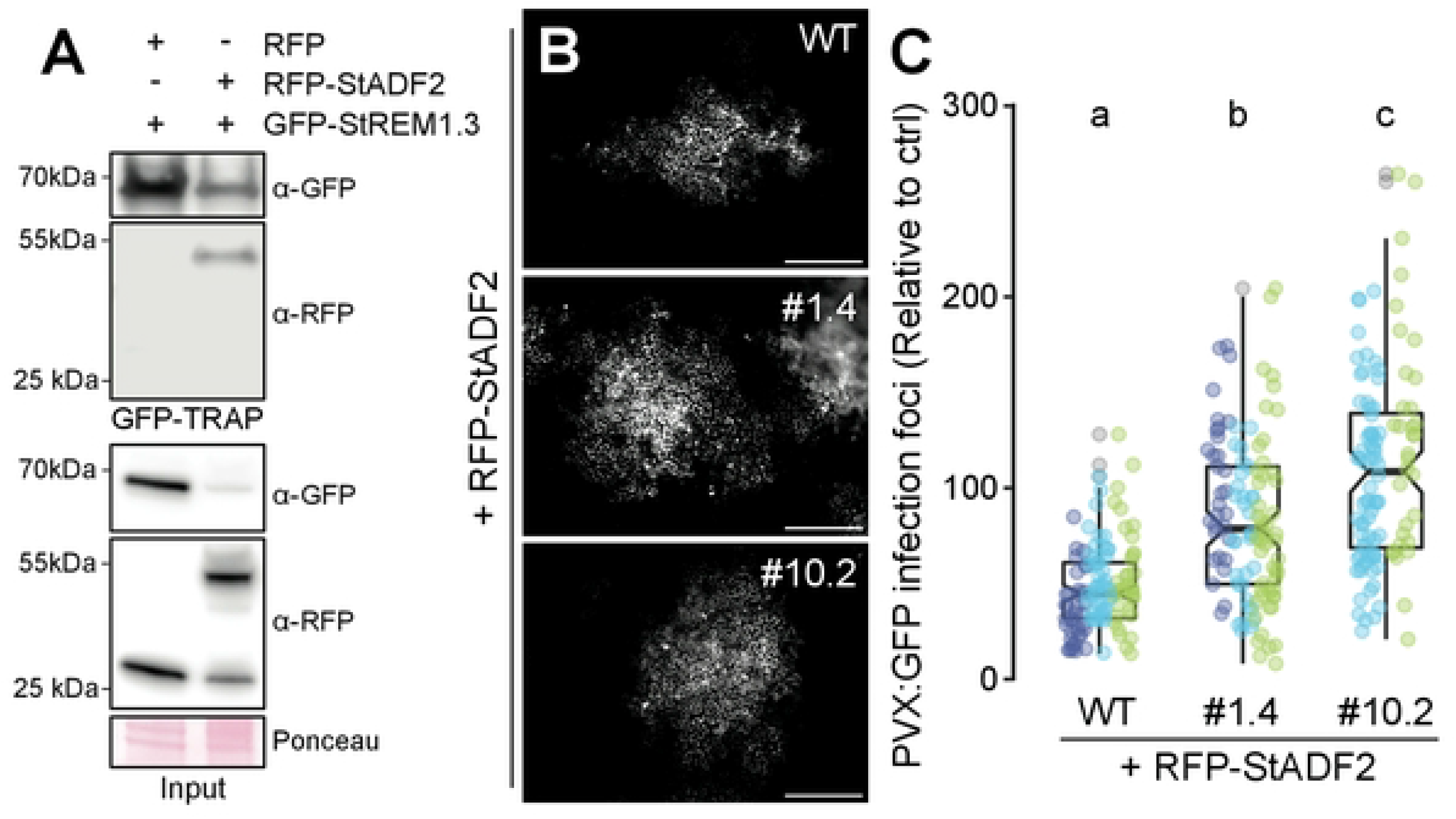
StADF2 limits PVX cell-to-cell movement in a REMORIN-dependent manner. **A**. Immunoprecipitation of GFP-StREM1.3 transiently co-expressed with either RFP or RFP-StADF2 in *N. benthamiana*. Blot stained with Ponceau is presented to show loading. Western blots were probed with α-GFP or α-RFP antibodies. **B-C**. PVX infection assays. Representative micrograph of PVX:GFP infection foci upon expression of RFP-StADF2 in WT and stable *REMORINs* knock-down transgenic *N. benthamiana* independent lines #1.4 and #10.2 (**B**) and corresponding quantification (**C**). PVX:GFP foci are normalized based on the average area observed in the absence of RFP-StADF2 overexpression for each genotype. Graphs are notched box plots, scattered data points show measurements, colors indicate independent experiments, number of infections foci n=114 for WT, n=113 for #1.4 and n=104 for #10.2. Conditions which do not share a letter are significantly different in Holm-Bonferroni’s multiple comparison test (p< 0.0001). Scale bar indicates 500 μm.

**Figure 5.**
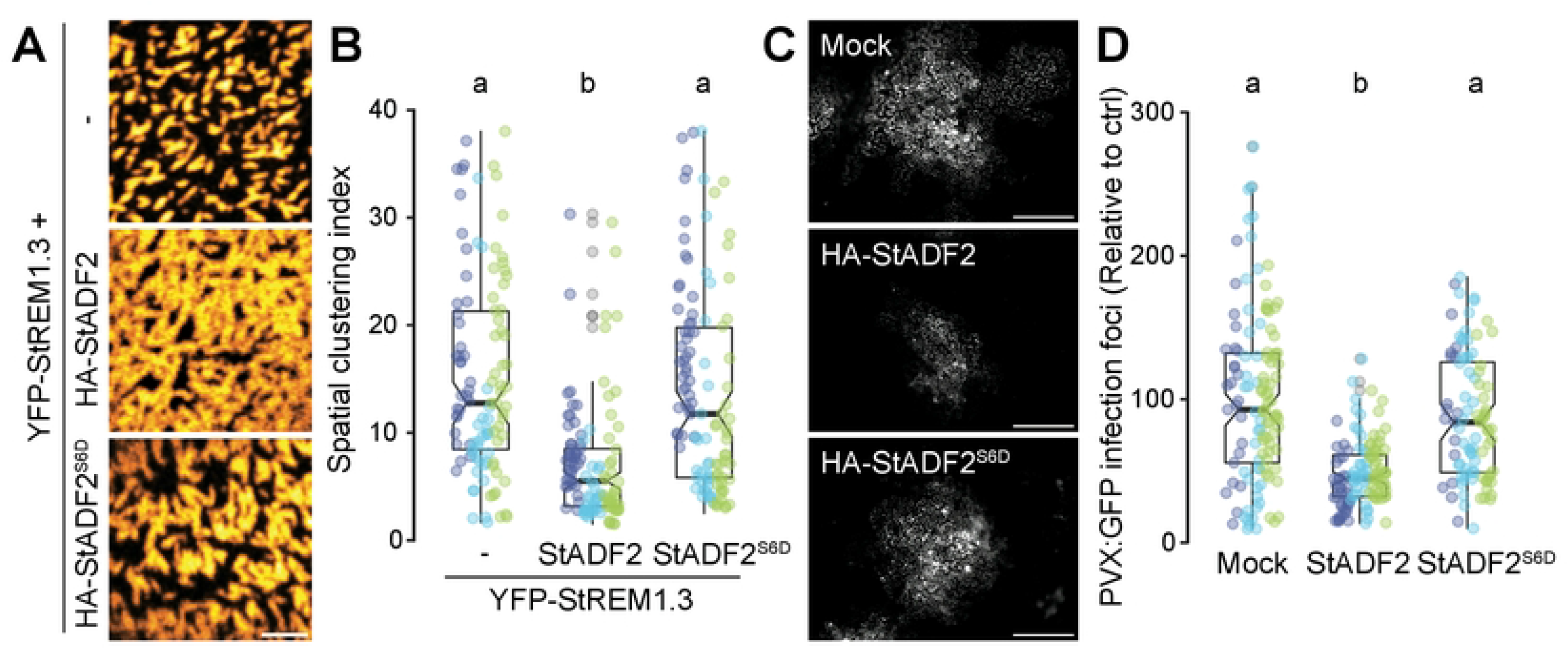
StADF2 alters StREM1.3 nanodomains to inhibit PVX cell-to-cell movement. **A-B**. Plasma membrane organization of YFP-StREM1.3 with or without co-expression of HA-StADF2 and HA-StADF2^S6D^. Representative confocal micrograph (**A**) and corresponding quantification of YFP-StREM1.3 spatial clustering index (**B**). Graphs are notched box plots, scattered data points show measurements, colors indicate independent experiments, n=38 cells for YFP-StREM1.3, n=38 cells for YFP-StREM1.3 + HA-StADF2 and n=38 cells for YFP-StREM1.3 + HA-StADF2^S6D^. Conditions which do not share a letter are significantly different in Holm-Bonferroni’s multiple comparison test (p< 0.0001). Scale bar indicates 2 μm. **C-D**. PVX infection assays. Representative micrograph of PVX:GFP infection foci upon expression of HA-StADF2 or HA-StADF2^S6D^ (**C**) and corresponding quantification (**D**). Graphs are notched box plots, scattered data points show measurements, colors indicate independent experiments, number of infections foci n=39 for mock, n=38 foci for HA-StADF2 and n=31 foci for HA-StADF2^S6D^. Conditions which do not share a letter are significantly different in Dunn’s multiple comparison test (p< 0.0001). Scale bar indicates 500 μm.

## DISCUSSION

How processes are regulated in space and time within the plasma membrane (PM) remains largely obscure. In plants, REMORINs are structural elements of the PM emerging as versatile regulatory components of various signaling pathways in plant development and plant-microbe interactions (Gui et al., 2016; Kohorn et al., 2016; Liang et al., 2018; Perraki et al., 2014). This functional versatility is suspected to be encoded in their variable and intrinsically disordered N-terminal domain (Gouguet et al., 2021). REMORINs IDDs have been found to be poly-phosphorylated in many plant species and across a wide range of physiological conditions (Gouguet et al., 2021). Here we show that the phosphorylation of StREM1.3 IDD modifies the chemical environment at the level of the protein backbone which may operate as a molecular switch in regulating its association with specific partners. In good agreement, we observed that a phosphomimic variant of StREM1.3 has both an increased ability to generate protein-protein interaction and a distinct interactome from the wild-type protein in yeast. These observations suggest the existence of context-dependent REMORINs phospho-codes and open way toward their identification and the study of their implication in the regulation of REMORINs-associated molecular complexes and signaling pathways.

Functional analysis of selected StREM1.3 interactors unveiled uncharacterized host regulators of PVX infection (Figure 3). These observations suggest that plant immune response to PVX involves the autophagy machinery (NbATG8i), water permeable channels (NbPIP1;3, NbTIP2;1), regulators of cellular redox status (NbTRX4) and iron-binding proteins (NbMT2B). Several orthologs of these proteins were previously linked to immunity in various plants species against pathogens of several kingdoms. For instance, the autophagy machinery has been found to play a central role in plant immunity against bacteria, fungi and viruses and can as well be manipulated by pathogens for their own benefits (Fu et al., 2018; Hafrén et al., 2018; Leary et al., 2018; Leong et al., 2022; F. Li et al., 2020; MacHaria et al., 2019; Niu et al., 2021; Üstün et al., 2018; Yang et al., 2018). Altogether, these observations suggest that REMORINs play a central role in regulating key cellular events in plant immunity, the definition of the precise mechanisms will require further investigation.

NbADF3 and StADF2 belong to the subclass 1 of the ADF protein family (Inada, 2017), which has been reported to be involved in multiple plant-pathogen interactions (Lu et al., 2015; Lu et al., 2020; Porter et al., 2012; Porter & Day, 2013; Tian et al., 2009). Subclass 1 ADFs have been identified in *Arabidopsis thaliana* plasmodesmata (PD) proteome (Brault et al., 2019), which may imply that their role in restricting of PVX propagation (Figure 3B and 4C) relies on a direct modulation of plasmodesmata. Further, actin disruption has been recently proposed as an immune response leading to callose deposition (Leontovyčová et al., 2020) and both AtADF4 and REMORINs are required for callose deposition in response to bacterial elicitors (Li et al., 2012; Ma et al., 2022). Since StREM1.3 regulates callose deposition at PD (Perraki et al., 2018) and that StADF2 limits PVX in a REMORIN-dependent manner (Figure 4C), modulation of actin cytoskeleton and of StREM1.3 ND organization by StADF2 may underly REMORIN-regulated callose accumulation at PD. Recently, evidences indicate that REMORINs and the actin cytoskeleton are working hand in hand during plant-microbe interactions. Arabidopsis REM1.2 promotes type-I Formin ND organization to foster actin nucleation in response to bacterial flagellin (Ma et al., 2022) while SYMREM1-induced membrane topology modifications during symbiosis depends on the actin cytoskeleton (Su et al., 2023). Our observations further reinforce the tight functional interplay between REMORINs and the actin cytoskeleton. Interestingly, two subclass 1 *Arabidopsis* ADFs (AtADF1 and AtADF4) as well as group 1 REM were shown to be phosphorylated by AtCPK3 (Dong & Hong, 2013a; Lu et al., 2020). Moreover, it was observed that CPK3 modulated actin cytoskeleton in response to bacterial elicitors (Lu et al., 2020) and limited PVX cell-to-cell movement in a REMORIN-dependent manner (Mehlmer et al., 2010; Perraki et al., 2018), which hints that CPK-ADF-REMs may correspond to a conserved immune signaling module against invading microbes. Stimuli-dependent organization of proteins in NDs is often presumed to be associated with initiation and activation of cellular processes. In good agreement, the stabilization of the small GTPase Rho-of-plants 6 (ROP6) into NDs upon auxin perception supports its function in regulating gravitropism (Platre et al., 2019). In addition, ROP6 forms specific domains promoting the production of reactive oxygen species upon osmotic stress (Smokvarska et al., 2020). However, recent reports suggest that the functional interplay between PM lateral organization and the functional status of its constituents is far more complex. While osmotic stimulation induces ND organization of the NADPH oxidase RBOHD and of ROP6 (Smokvarska et al., 2020), it concomitantly leads to an increased mobility of the aquaporin PIP2;1 and of the proton pump ATPase AHA2 (Martinière et al., 2019). In yeast, the arginine permease Can1 accumulates into so-called membrane compartment occupied by Can1 (MCC) in its inactive, substrate-free form, while its active form shows a dispersed organization within the PM (Gournas et al., 2018). Similarly, substrate perception by the yeast methionine permease Mup1 induces its reorganization from NDs to a disperse plasma membrane network (Busto et al., 2018). We previously showed that native ND organization of StREM1.3 is required to support its function in limiting PVX cell-to-cell movement (Gronnier et al., 2017). Yet, active phosphomimic form of StREM1.3 presents a dispersed organization (Perraki et al., 2018; Figure 1A) suggesting that StREM1.3 is activated within NDs and subsequently dispersed. Here, we show that StADF2 alters StREM1.3 ND organization and limits PVX cell-to-cell movement in a REMORIN-dependent manner (Figure 4). Mutating a conserved single residue reported to affect ADFs affinity to actin inhibits StADF2 effect on StREM1.3 ND organization and PVX cell-to-cell movement (Figure 5). Altogether, these observations suggest that active modulation of actin cytoskeleton by StADF2 changes StREM1.3 nanoscale organization to limit PVX cell-to-cell movement. In mammals, actin depolymerization have been shown to promote B cell activation by enabling B cell antigen receptor clustering in the immune synapse (Droubi et al., 2022; Mattila et al., 2016; Wang & Huse, 2022). Since REMORINs have been proposed to regulate various cell-surface receptor signaling pathways (Bücherl et al., 2017; Gui et al., 2016; Liang et al., 2018), a similar mechanism could apply in response to PVX and in other contexts. Our study shades light on the activation of a PM-based immune response by membrane scaffold dispersion. We envision that analogous mechanisms may be conserved across organisms.

## MATERIALS AND METHODS

### Plant materials, culture and transformation

*N. benthamiana* plants were cultivated in controlled conditions (16 h photoperiod, 25 °C). *N. benthamiana* plants were transiently transformed using the *Agrobacterium tumefaciens* strain GV3101 as previously described (Gronnier et al., 2017). For subcellular localization study and western blotting analyses, plants were analyzed 2 days after infiltration (dai). For PVX:GFP spreading assays, plants were observed 5 dai. The *N. benthamiana* transgenic RNAi knock-down lines hpREM #1.4 and hpREM #10.2 were previously described (Perraki et al., 2018).

### PVX cell-to-cell movement

The assays were conducted as previously described (Perraki et al., 2018). Agrobacteria solution (OD^600nm^ = 0.001) carrying PVX:GFP and the helper plasmid pSoup was mixed to equal volume with agrobacteria carrying a binary plasmid encoding for tested proteins (OD^600nm^ = 0.2) and co-infiltrated in leaves of 3 weeks old *N. benthamiana*. Plants were observed at 5 dai using Zeiss Axiozoom V16 equipped with a UV lamp, an excitation filter (450-490 nm) and an emission filter (500-550 nm) to detect GFP signal. Foci were automatically analyzed using the Fiji software (Schindelin et al., 2012) via a homemade macro.

### Membrane-based split-ubiquitin Yeast two-hybrid

Split ubiquitin assays were performed using the yeast two-hybrid system from DUAL membrane system (Dual systems Biotech AG). StREM1.3^WT^ and StREM1.3^S74D T86D S91D^ coding sequences were amplified by PCR using SfiI restriction site–containing primers (Supplemental Table 2) and subsequently cloned in pBT3N bait and pPR3N prey plasmids. To test the functionality of StREM1.3 in this system, THY.AP4 cells were sequentially transformed with pBT3N:StREM1.3 and pPR3N:StREM1.3 or pPR3N empty vector as a negative control. The cDNA library was constructed following the manufacturer’s instructions (Evrogen) using approximatively 700 ng total RNA of epidermis peeled from healthy and PVX-infected *N. benthamiana* leaves, harvested three days after agroinfiltration. For screening, NubG-cDNA library was used for transformation of THY.AP4 yeast strain *MATa, ura3-, leu2-, lexA-lacZ∷TRP1, lexA∷HIS3, lexA∷ADE2*) previously transformed with pBT3N:StREM1.3 or pBT3N:StREM1.3^DDD^. Positive clones were selected on synthetic medium (SD) lacking adenine, histidine, tryptophan and leucine (-AHWL) and subsequently tested for β-galactosidase activity. To measure β-galactosidase activity, yeasts were grown on SD-TL for two days at 28°C. Plates were then covered with a X-Gal-agarose buffer (0.5% agarose, 0.5 M phosphate buffer, pH 7.0, 0.002% X-Gal) and incubated at 37°C for 10 to 20 min. To identify the proteins expressed in Yeast positive clones obtained from the SUY2H exploratory screens, the corresponding plasmids were isolated from Yeast, subsequently propagated in *E. coli*, isolated and analyzed by Sanger sequencing. Individual coding sequences were BLAST against *N. benthamiana* Genome v1.0.1 predicted cDNA (https://solgenomics.net/).

### *In silico* analyses

During SUY2H exploratory screens, identification of proteins expressed in yeast positive clones was retrieved using BLASTn algorithm against *N. benthamiana* Genome v1.0.1 predicted cDNA. Closest orthologs of the identified StREM1.3-interacting proteins in *A. thaliana* were retrieved on The Arabidopsis Information Resource TAIR using BLASTp algorithm. The 500 hundreds closest homologs of SlADF2 were retrieved from 101 plant species in Phytozome (Goodstein et al., 2012) using BLASTp. Protein alignment was computed using MUltiple Sequence Comparison by Log-Expectation (MUSCLE; (Edgar, 2004)) using BLOSUM62 matrix, an -sv profile scoring method with following parameters: Anchor spacing:32, diagonal break:1, diagonal length:24, diagonal margin:5, gap extension penalty:-1, gap open penalty:-12, hydro:5 and hydro factor1.2, through the JABAWS server (Troshin et al., 2011). Among 500 ADF protein sequences, 41 are predicted splice variants lacking the first 7 amino acids and were excluded for sequence logo analysis, generated using WebLogo (Crooks et al., 2004).

### Molecular cloning

Candidate genes selected from the screen were cloned from the cDNA bank generated for the SUY2H screen using primers designed to amplify full length coding sequences. All vector constructs were generated using classical Gateway cloning strategies (www.lifetechnologies.com), using pDONR211 and pDONR207 as entry vectors and pK7WGY2, pK7YWG2, pK7WGR2, pK7RWG2 (Karimi et al., 2002), pGWB14, pGWB15 (Nakagawa et al., 2007) and pSite BiFC as destination vectors (Martin et al., 2009). StADF2 and StADF2^S6D^ bearing attL sequences were synthesized (https://www.genscript.com/) in a pUC57 vector to be cloned in aforementioned destination vectors. The StREM1.3^S74D T86D S91D^ was previously described (Perraki et al., 2018). For GFP-StREM1.3 expression in *Saccharomyces cerevisiae*, StREM1.3^WT^ and StREM1.3^DDD^ coding sequences were PCR amplified with oligonucleotides listed in supplemental table S2, digested by BamHI–NsiI and cloned at BamHI–PstI sites of the plasmids p2717 (pCM189 modified by the introduction of a myc epitope tag downstream of the tet promoter (Escusa et al., 2006)). The resulting plasmids (respectively pMC101 and pMC104) were transformed into a wild type strain BY4741 (*MATa, his3Δ1, leu2Δ0, met15Δ0, ura3Δ0*). All plasmids were propagated using the NEB10 *E. coli* strain (New England Biolabs).

### Protein expression in E. coli and purification

StREM1.3^1-116^ was obtained as previously described (Legrand et al., 2022). Shortly, StREM1.3^1-116^ was expressed in BL21-DE3 cells in minimal medium with ^13^C-labelled glucose and ^15^NH^4^Cl, by addition of 1 mM IPTG at OD^600^ = 0.6-0.8 and incubation at 37°C for 3h. Cells were lysed by sonication and the supernatant was loaded onto a HisTrap column (GE Healtchare) equilibrated in 20 mM HEPES 150 mM NaCl, 20 mM imidazole, 0.02% NaN_3_, pH=7.4 and eluted with 20 mM HEPES, 150 mM NaCl, 500 mM imidazole, 0.02% NaN_3_, pH=7.4. For TEV cleavage, eluted StREM1.3^1-116^ was adjusted to 1 mM DTT and 0.5 mM EDTA, the TEV protease was added in a ~1/200 TEV/REM^1-116^ mass ratio and the reaction was incubated for 3h at room temperature then desalted against 10 mM HEPES, 50 mM NaCl, 0.02% NaN_3_, pH=7.5 with a HiPrep column (GE Healthcare). Under native conditions, StREM1.3^1-116^ was loaded onto a Histrap column equilibrated with 20 mM HEPES, 150 mM NaCl, 0.02% NaN_3_, pH=7.4 and eluted with the same elution buffer as above. Under denaturing conditions, this step was performed in buffers containing 7M urea. Finally, StREM1.3^1-116^ was desalted again against 10 mM HEPES, 50 mM NaCl 0.02%, NaN_3_ pH=7.5. AtCPK3-GST recombinant protein was produced in BL21-DE3-pLys and purified as previously reported (Legrand, et al., 2022). As a negative control, AtCPK3-GST was inactivated by heating at 95°C for 10 min.

### *In vitro* phosphorylation

*In vitro* phosphorylation of 0.5 mM of StREM1.3^1-116^ by AtCPK3 was done as previously published (Legrand et al., 2022). For NMR analysis, the StREM1.3^1-116^ sample was adjusted to 10 mM MgCl_2_, 1 mM CaCl_2_, 1 mM DTT and 3 mM ATP. The reaction was initiated by the addition of 88 μM of AtCPK3 and incubation at 20°C.

### Liquid-state Nuclear magnetic resonance (NMR) spectroscopy

NMR spectroscopy was performed on a Bruker Advance NEO spectrometer operating at 700 MHz for proton with a TXI 5 mm probe. The samples were adjusted to 9/1 H_2_O/D_2_O (v/v) to lock the magnetic field. Monitoring of phosphorylation kinetics was performed using a ^1^H-^15^N HMQC pulse sequence (Schanda & Brutscher, 2005), with 2 scans per timepoint (*i.e* 1 min 42 s to acquire one spectrum). The spectra were processed and analyzed using TopSpin (Bruker).

### Confocal laser scanning microscopy and image analysis

Live imaging was performed using a Zeiss LSM 880 confocal laser scanning microscopy system using a 68x objective and the AiryScan detector. YFP fluorescence was observed using an excitation wavelength of 488 nm and an emission wavelength of 453nm. RFP and mCherry fluorescence were observed using an excitation wavelength of 561 nm and detected at 579 nm. Acquisition parameters were kept identical across experiments. The SCI was calculated as previously described (Gronnier et al., 2017). Briefly, 10 μm lines were plotted across the samples and the SCI was calculated by dividing the mean of the 5 % highest values by the mean of 5 % lowest values. Three lines were randomly plotted per cell.

To disrupt actin cytoskeleton integrity, Cytochalasin D (40 μg/mL) dissolved in DMSO was infiltrated 24 hours before observation, infiltration of DMSO for 24 hours was used as corresponding mock control.

Yeast cells were observed on a fully-automated Zeiss 200M inverted microscope (Carl Zeiss, Thornwood, NY, USA) equipped with an MS-2000 stage (Applied Scientific Instrumentation, Eugene, OR, USA), a Lambda LS 300-Watt xenon light source (Sutter, Novato, CA, USA), and a 100 × 1.4 NA Plan-Apochromat objective. GFP imaging was done using a FITC filter (excitation, HQ487/25; emission, HQ535/40; beam splitter, Q505lp). Images were acquired using a Prime sCMOS 95B camera (Photometrics, Tucson, AZ, USA). The microscope, camera, and shutters (Uniblitz, Rochester, NY, USA) were controlled by SlideBook software 5. 0. (Intelligent Imaging Innovations, Denver, CO, USA).

### Western-blotting

Proteins were extracted from *N. benthamiana* leaf tissue transiently expressing HA-tagged proteins in 100 mM Tris (pH 7.5) containing 3% SDS, 5mM EDTA and 2% Protease inhibitor (Roche, Complete) boiled for 10 min at 70°C in SDS loading buffer, and cleared by centrifugation. The protein extracts were then separated by SDS-PAGE, transferred to polyvinylidene difluoride (PVDF) membranes (BioRad), blocked with 5% skimmed milk in TBS-Tween 0.05%, and incubated with an anti-HA antibody coupled to horse radish peroxidase (Roche). The resulting western-blots were developed using an ECL Prime Kit (GE Healthcare) and detected with an ImageQuant 800 (Amersham).

### Co-immunoprecipitation

Immunoprecipitation assays were performed as previously described in (Dagdas et al., 2016) with minor modifications. Approximately 3 g of *N. benthamiana* leaves were ground with a mortar and pestle in liquid nitrogen and homogenized in 6 mL of extraction buffer (50 mM Tris- HCl, pH 7.5, 150 mM NaCl, 10% glycerol, 5 mM DTT, 1 mM EDTA, 2% (w/v) polyvinylpolypyrrolidone, 1% (v/v) COmplete protease inhibitor cocktail [Roche], 0.1% (v/v) IGEPAL). Samples were then centrifuged for 20 min with 2000 x g at 4°C. Immunoprecipitation was performed by adding 30 μl of GFP-Trap coupled to agarose beads (ChromoTek) and samples were continuously agitated for 2 hours at 4°C. Beads were subsequently washed five times with extraction buffer and eluted with 30 μl of 2X Laemmli buffer at 70 °C for 10 minutes.

### Statistical analyses

Statistical analyses were carried out using Prism 6.0 software (GraphPad). As mentioned in the figure legends, statistical significances were assessed using non-parametric Kruskal-Wallis bilateral tests combined with post-hoc Dunn’s multiple pairwise comparisons, or using a two-way non-parametric Mann-Whitney test.

### Accession numbers

NbADF3 (NbS00025994g0001.1), NbCRT3 (NbS00018258g0010.1), NbMT2B (NbC25904295g0003.1), NbATG8I (NbS00005942g0011.1), NbPIP1C (NbS00006841g0003.1), NbTIP2;1, (NbS00006781g0007.1), NbDRM1 (NbS00004204g0005.1), NbGST8 (NbS00007668g0012.1), NbTRXH4 (NbS00049748g0003.1), NbRabA4A (NbS00057294g0003.1), StREM1.3 (NP_001274989), StADF2 (Soltu.DM.04G007350).

#### ACKNOWLEDGEMENTS

We thank the Bordeaux Imaging Center, part of the National Infrastructure France-BioImaging supported by the French National Research Agency (ANR-10-INBS-04). This work was supported by the French National Research Agency (grant no. ANR-19-CE13-0021 to SGR, SM, VG and doctoral fellowships to MDJ, PG and JG), and the German Research Foundation (DFG) grant CRC1101-A09 to JG, the IPS2 benefits from the support of the LabEx Saclay Plant Sciences-SPS (ANR-10-LABX-0040-SPS). We thank Isabel Monte and members of the NanoSignaling Lab for discussions and critical reading of the manuscript.

The authors declare no conflict of interest

#### AUTHORS CONTRIBUTION

ALe purified labelled StREM1.3^1-116^ and performed NMR experiments

ALe, BH and AL analyzed NMR results

JG and VG built the cDNA libraries

AML and IS localized fluorescent tagged StREM1.3 in yeast

JG performed the split-ubiquitin assays

PG, MDJ and JG performed and analyzed virus propagation

MDJ performed confocal microscopy

MB performed in vitro kinase assay

JG, MD, BH, AL, VG, SM designed the project

MDJ and JG assembled the figures

MDJ and JG wrote the manuscript with input from all authors.

---

**Figure supplemental 1.**
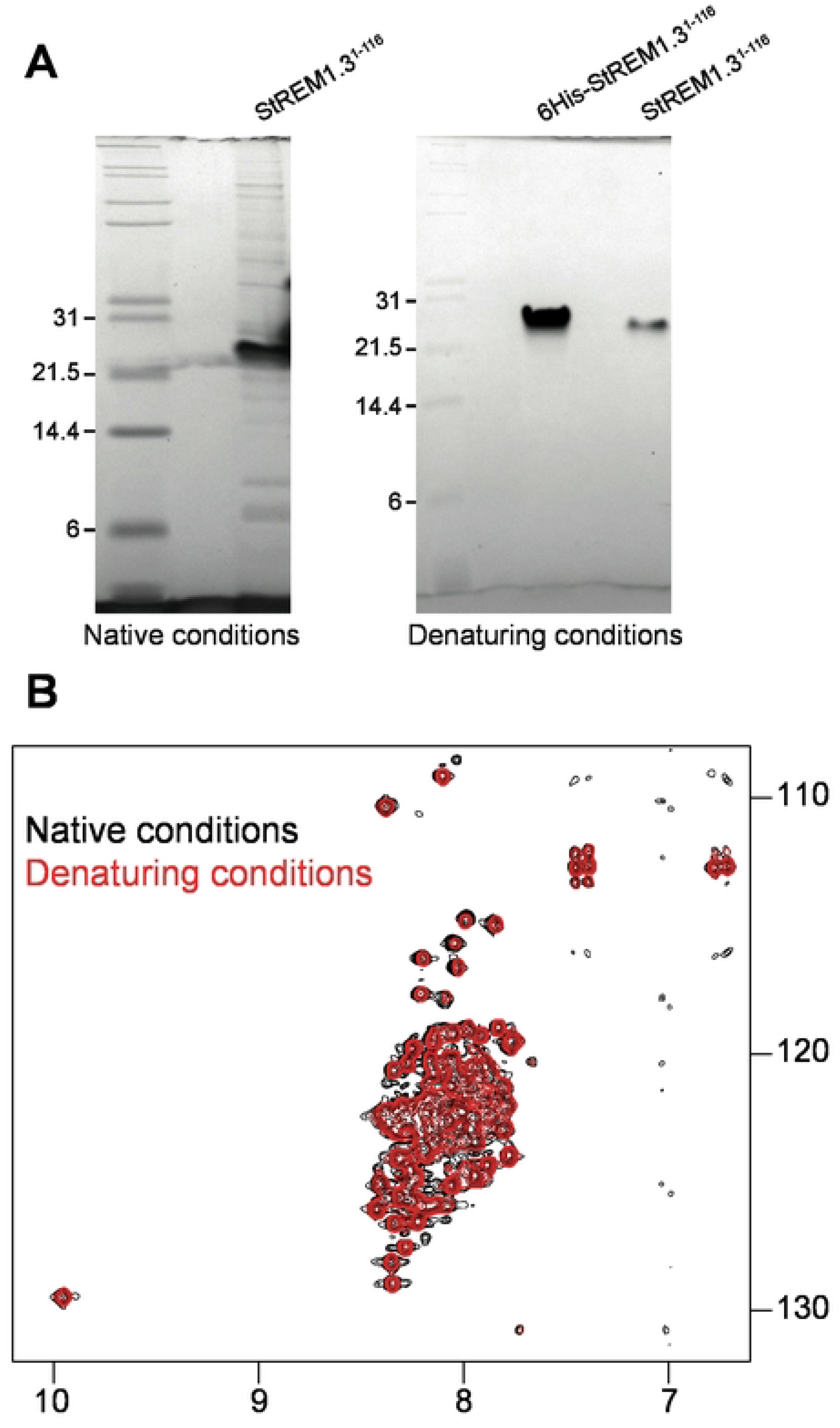
StREM1.3 N-terminal domain is intrinsically disordered. **A**. SDS-PAGE of purified untagged StREM1.3^1-116^ under native and denaturing conditions. **B**. ^1^H-^15^N HMQC spectra of REM^1-116^ purified under native (black) or denaturing conditions using 7M of urea (red). Temperature: 10°C.

**Figure supplemental 2.**
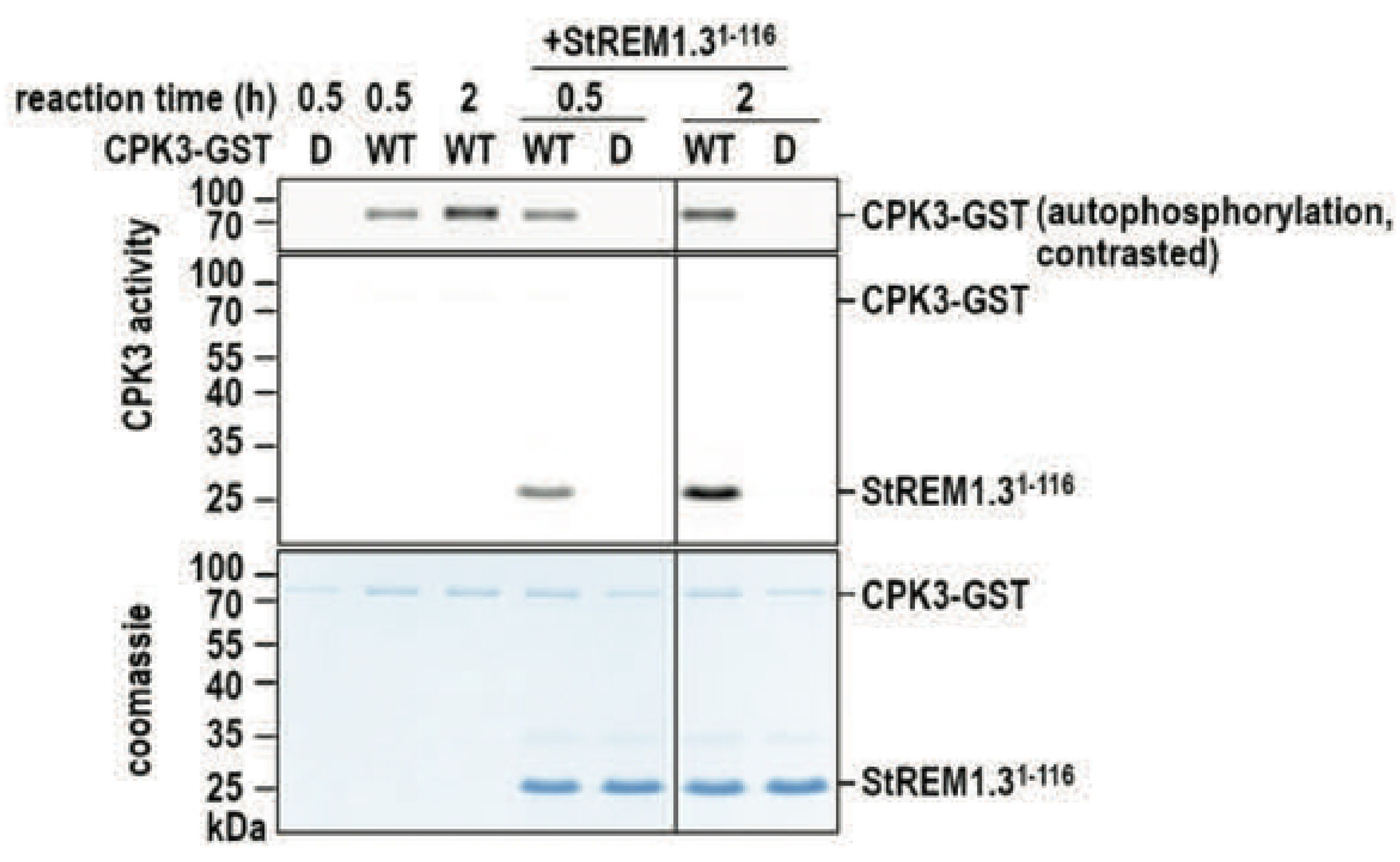
AtCPK3 phosphorylates StREM1.3^1-116^ under NMR conditions. **B**. *In vitro* kinase assay to confirm phosphorylation of StREM1.3^1-116^ under liquid-state NMR conditions. StREM1.3^1-116^ incubated for either 30 min or 2 h with wild-type AtCPK3-GST untreated (WT) or heat-inactivated (D).

**Figure supplemental 3.**
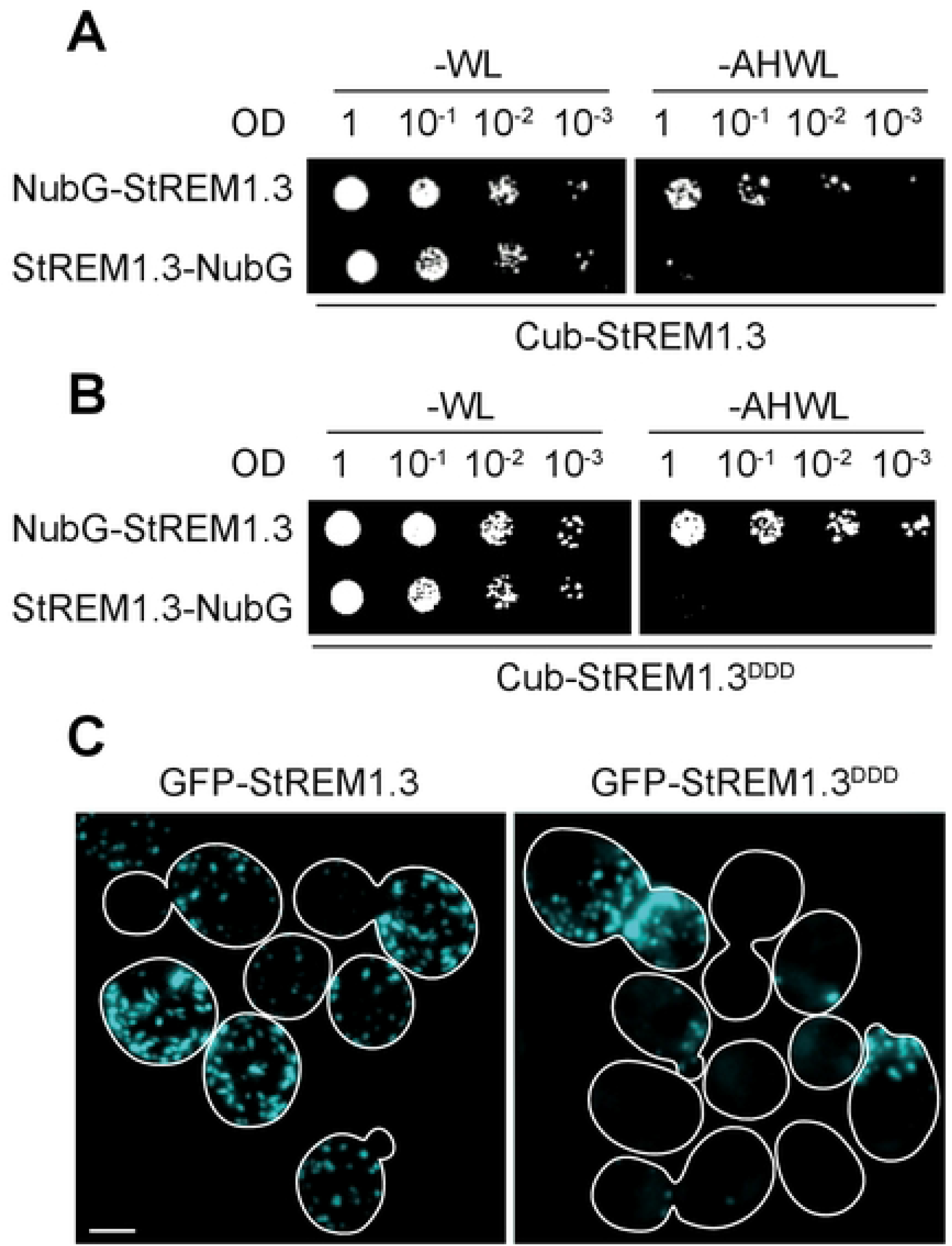
Cub-StREM1.3 and Cub-StREM1.3^DDD^ are functional baits in SUY2H. **A-B**. Drop-tests of SUY2H assays testing binary interactions between Cub-StREM1.3 (**A**) with NubG-StREM1.3 or StREM1.3-NubG, or Cub-StREM1.3^DDD^ (**B**) with NubG-StREM1.3 or StREM1.3-NubG. **C**. Representative micrograph of GFP-StREM1.3 and GFP-StREM1.3^DDD^ imaged in proliferating *S. cerevisiae*. Scale bar indicates 2 μm.

**Figure supplemental 4.**
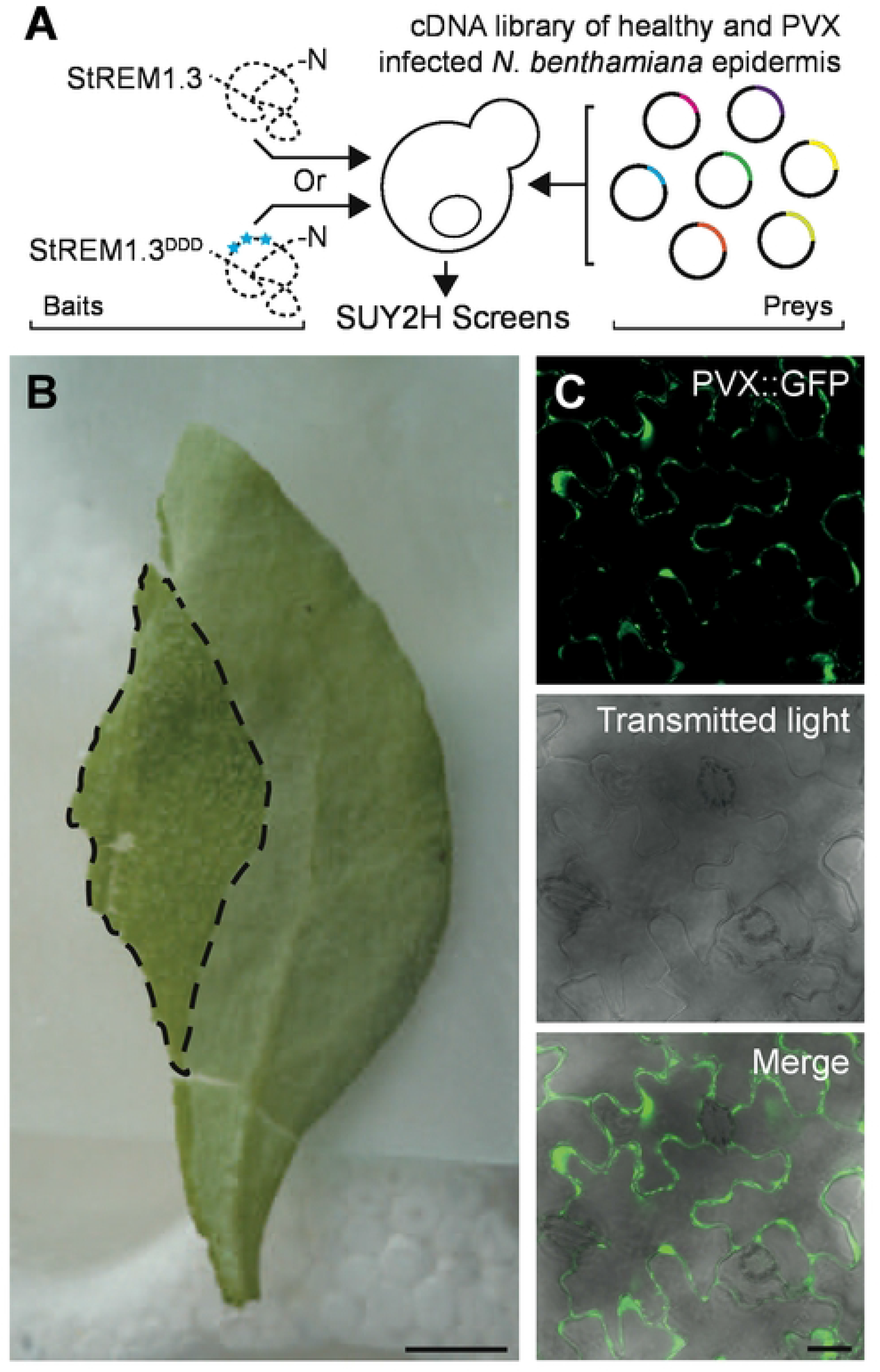
Rational design of the SUY2H screens. **A**. Schematics depicting the experimental design. Cub-StREM1.3 and Cub-StREM1.3^DDD^ were used as baits against a cDNA library obtained from PVX-infected *N. benthamiana* epidermis in side-by-side screens. **B**. Picture of a *N. benthamiana* leaf peeled. Scale bar indicates 5 cm. **C**. Confocal micrograph of peeled *N. benthamiana* epidermis infected with PVX:GFP. Scale bar indicates 5 μm.

**Figure supplemental 5.**
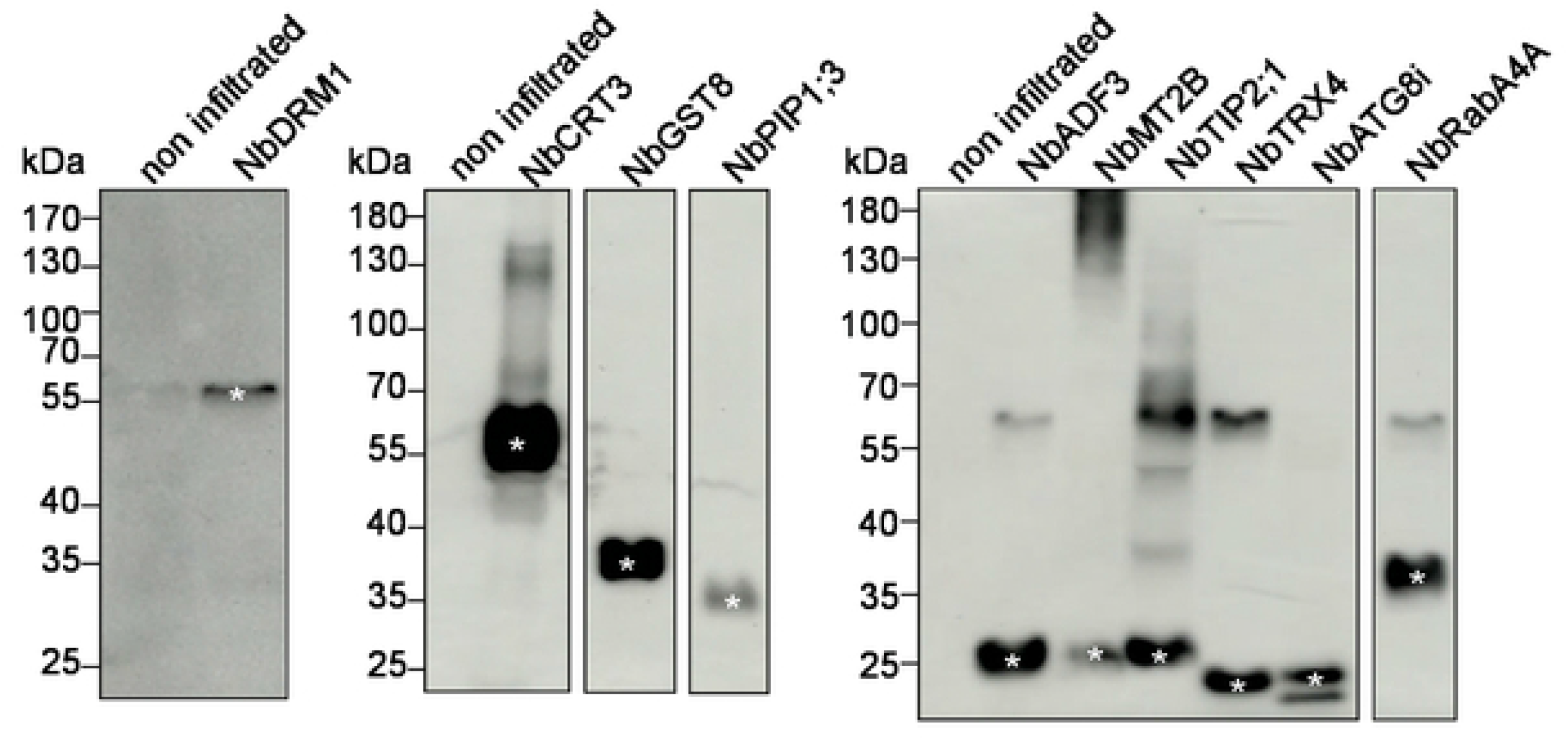
Western blots analyses of the expression of protein candidates in PVX cell-to-cell movement assays. Protein extracts from *N. benthamiana* leaves expressing HA-tagged candidates were analyzed by western blots with α-HA antibody. In case of multiple bands, a white star points out the band corresponding to the expected molecular weight. Abbreviations are as followed: Dormancy-associated Protein-like 1 (NbDRM1), Calreticulin 3 (NbCRT3), Glutathione S-transferase 8 (NbGST8), Plasma membrane intrinsic protein 1C (NbPIP1C), Actin Depolymerizing Factor 3 (NbADF3), Metallothionein 2B (NbMT2B), Delta tonoplast integral protein (NbTIP2;1), Thioredoxin superfamily protein 4 (NbTRX4), Autophagy 8I (NbATG8i), GTPase protein (NbRabA4A).

**Figure supplemental 6.**
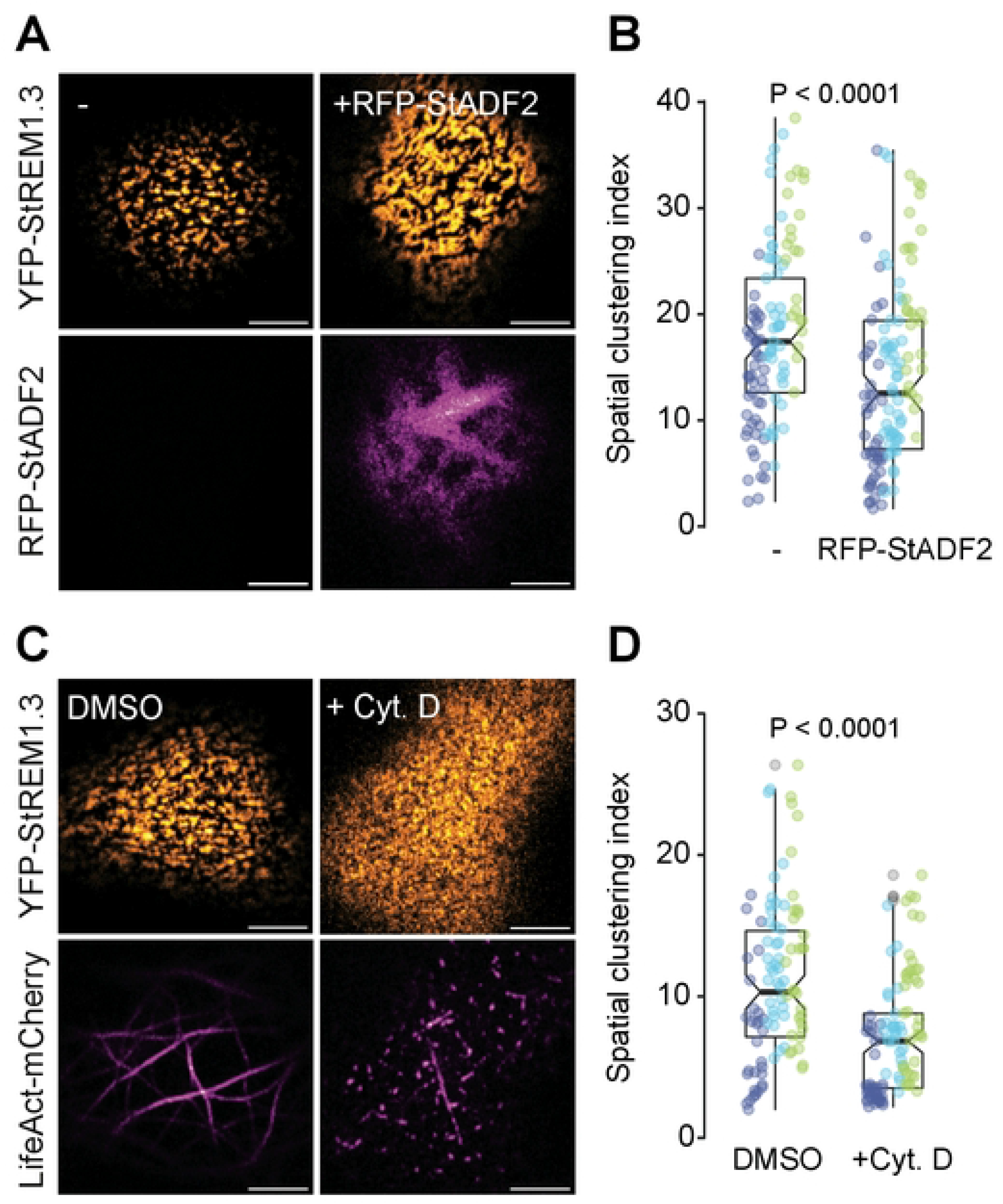
StADF2 affects YFP-StREM1.3 nanodomain organization and YFP-StREM1.3 nanodomain relies on the actin cytoskeleton integrity. **A-B**. Plasma membrane organization of YFP-StREM1.3 with or without co-expression of RFP-StADF2. Representative confocal micrograph (**A**) and corresponding quantification of YFP-StREM1.3 spatial clustering index (**B**) are shown. Graphs are notched box plots, scattered data points show measurements, colors indicate independent experiments, n=39 cells for YFP-StREM1.3, n=41 cells for YFP-StREM1.3 + RFP-StADF2. P values report two-tailed non-parametric Mann-Whitney test. Scale bar indicates 5μm. **C-D**. Plasma membrane organization of YFP-StREM1.3 co-expressed with LifeAct-mCherry to label actin upon 80 μM Cytochalasin D treatment for 24h or corresponding mock control (DMSO). Representative confocal micrograph (**C**) and corresponding quantification of YFP-StREM1.3 spatial clustering index (**D**) are shown. Graphs are notched box plots, scattered data points show measurements, colors indicate independent experiments, n=34 cells for mock treatment, n=34 cells for Cytochalasin D treatment. P values report two-tailed non-parametric Mann-Whitney test. Scale bar indicates 5μm.

**Figure supplemental 7.**
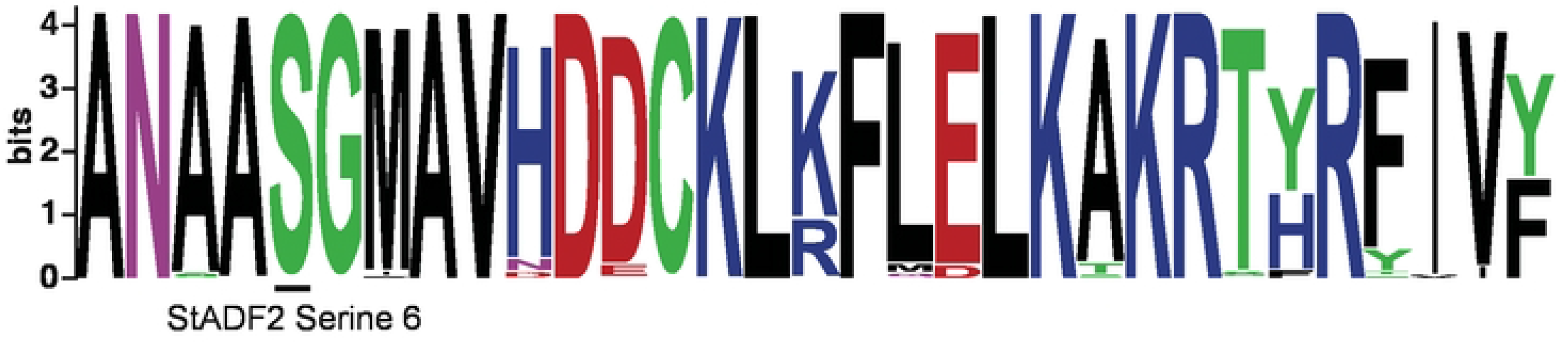
Serine 6 is conserved among ADFs. Sequence logo of ADFs N-termini generated from 459 protein sequences retrieved from 101 plant species using BLASTp and StADF2 as query.

